# Relative rDNA copy number is not associated with resistance training-induced skeletal muscle hypertrophy and does not affect myotube anabolism *in vitro*

**DOI:** 10.1101/2024.03.04.583417

**Authors:** Joshua S. Godwin, J. Max Michel, Andrew T. Ludlow, Andrew D. Frugé, C. Brooks Mobley, Gustavo A. Nader, Michael D. Roberts

**Author notes:** Address correspondence: Michael D. Roberts, PhD, Auburn University Alumni Professor, School of Kinesiology, Auburn University, 301 Wire Road, Auburn, AL 36849.

## Abstract

Ribosomal DNA (rDNA) copies are organized in tandem repeats across multiple chromosomes, and inter-individual variation in rDNA copy number has been speculated to be a modifier of the hypertrophic responses to resistance training. In the current study, 82 apparently healthy participants (n=53 males, 21±1 years old; n=29 females, 21±2 years old) performed 10-12 weeks of supervised full-body resistance training. Whole-body, mid-thigh, and histological skeletal muscle hypertrophy outcomes were determined, as was relative rDNA copy number from pre-intervention vastus lateralis (VL) biopsies. Pre- and post-intervention VL biopsy mRNA/rRNA markers of ribosome content and biogenesis were assayed in all participants, and these targets were also assayed in the 29 females 24 hours following their first workout bout. Across all 82 participants, no significant associations were evident between relative rDNA copy number and training-induced changes in whole body lean mass (r = -0.034, p=0.764), vastus lateralis thickness (r = 0.093, p=0.408), mean myofiber cross-sectional area (r = -0.128, p=0.259), or changes in muscle RNA concentrations (r = 0.026, p=0.818). Several significant, positive associations in females support ribosome biogenesis being linked to training-induced myofiber hypertrophy. Follow-up studies using LHCN-M2 myotubes demonstrate a reduction in relative rDNA copy number induced by bisphenol A (BPA). However, BPA did not significantly affect myotube diameter or prevent insulin-like-growth factor-induced hypertrophy. These findings provide strong evidence that relative rDNA copy number is not associated with myofiber anabolism and provide further mechanistic evidence for ribosome biogenesis being involved in this phenomenon.

## INTRODUCTION

Skeletal muscle can molecularly adapt to a multitude of internal and external stressors. In the case of skeletal muscle hypertrophy in response to mechanical overload, an increase in protein accumulation/synthesis is required to facilitate the growth of myofibers [1-4]. Ribosomes are the macromolecules responsible for facilitating protein translation, thus regulating protein synthesis. To meet the demand for increased protein synthesis during hypertrophic stimuli, ribosomal capacity also must increase [1]. Protein synthesis, with respect to ribosomes, is regulated via changes in ribosomal content (translational capacity) and/or ribosomal activity (translational efficiency) [5-7]. Changes in ribosomal content during resistance training occurs via *de novo* ribosome biogenesis, which requires coordination of all three RNA polymerases (RNA Pol-I, II, and III) but is primarily driven by transcription of the 45S rDNA genes via Pol-I [8, 9]. Pol-I-mediated rDNA transcription results in the 47/45S pre-rRNA transcript that is subsequently processed into the mature 28S, 18S, and 5.8S rRNAs [10]. Pol-I activity is supported by several factors including upstreaming binding factor (UBF, also known as UBF1), the selectivity factor-1 (SL-1), transcription initiation factor-1 (TIF-1A), and other accessory proteins; collectively, these supporting factors are deemed the “Pol-I regulon” [11-15]. Furthermore, the transcription of rDNA into the 45S pre-rRNA is considered the rate-limiting step for ribosome biogenesis [5, 16, 17].

An increase in ribosomal capacity has been reported following both acute bouts [16, 18-20] and longer-term periods [19, 21-23] of resistance training in humans. Moreover, Nakada et al. [24] utilized a rodent model of incremental surgical overload to show that a greater accumulation of rRNA levels (i.e., ribosomal content) was associated with higher degrees of skeletal muscle hypertrophy. At the onset of skeletal muscle hypertrophy, transcription of rDNA genes occurs in a pulsative fashion peaking from 24-48 hours resulting in the accumulation of rRNAs. rDNA transcription is further regulated epigenetically via methylation status and various histone modifications [16, 25-28].

Another unique aspect that may influence rDNA transcription is the inter-individual variability of rDNA copy number. Copies of the rDNA gene exist in tandem repeats spread across multiple chromosomes [29, 30]. However, while relative rDNA copy number has been linked to enhanced ribosome biogenesis 24 hours following a resistance exercise bout in humans [16], no human study has sought to investigate the association between this genetic feature and resistance training-induced skeletal muscle hypertrophy. Therefore, the primary purpose of the current investigation was to determine if relative rDNA copy number was associated with changes in hypertrophic outcomes in humans. Moreover, we sought to examine if relative rDNA copy number was associated with changes in markers of ribosome biogenesis following one bout as well as chronic resistance training. We hypothesized that relative rDNA copy number would be positively associated with changes in various markers of skeletal muscle hypertrophy following chronic resistance training. Moreover, we hypothesized that relative rDNA copy number would be positively associated with increases in ribosome biogenesis markers following acute and chronic resistance training.

## METHODS

### Participants and ethical approvals

Human specimens were from apparently healthy male (n=53, 21±1 years old, 77.5±10.6 kg, 24.1±2.9 kg/m^2^) and female (n=29; 21±2 years old, 67.5±11.0 kg, 23.4±3.4 kg/m^2^) participants from two prior studies performed in our laboratory [31, 32]. Study procedures for both studies were approved by the Auburn University Institutional Review Board (male study approved protocol #: 15-320, female study approved protocol #: 19-249) and conformed to the latest standards of the Declaration of Helsinki. All participants were fully aware of the study procedures prior to providing verbal and written consent. Additional details concerning inclusion and exclusion criteria, as well as details related to the procedures have been previously reported [31, 32].

### General Training Protocol

Female participants completed a progressive resistance training regime consisting of 2 days per week for 10 weeks. Briefly, subjects performed leg press, bench press, leg extension, hex-bar deadlift, and cable pulldowns for 4 sets of 10 repetitions (day 1) and 5 sets of 6 repetitions (day 2). Load was increased weekly by ∼5% for higher volume days (day 1) and ∼9% for higher load days (day 2). Additionally, load could be adjusted for each exercise depending on the rating of perceived exertion provided by participants (RPE). All training was conducted under the supervision of research staff.

Male participants completed a different protocol comprised of 3 days per week of daily undulating periodization resistance training for 12 weeks. Briefly, subjects performed barbell back squats, bench press and bent-over rows for 4 sets of 10 repetitions (day 1), 6 sets of 4 repetitions (day 2), and 5 sets of 6 repetitions (day 3). Training load was increased weekly by ∼5-10% for each day and was further adjusted based on RPE. Again, all training was conducted under the direct supervision of research staff.

### Acute training paradigm in female participants

The first/naïve training bout occurred in conjunction with assessment of three-repetition maximum (3-RM) strength testing. Participants completed a general warm-up followed by 3-RM testing for 45° leg press, barbell bench press, and hex-bar deadlift. Following warm-up sets at 10 and 5 repetitions, load was incrementally increased by 2-3% until 3-RM was achieved. The 3-RM value was then used to calculate an estimated 1-RM. Following completion of 3-RM strength testing, participants completed 2 additional sets of 10 repetitions with 50% of the estimated 1-RM on leg press, bench press, and hex-bar deadlift. The overall training session included 3-RM testing and 2 subsequent sets at 50% est. 1-RM for all exercises.

### Testing sessions

#### Urine Specific Gravity

Prior to the first and last testing sessions, participants donated a urine sample (∼5 mL). Samples were analyzed using a handheld refractometer (ATAGO) for urine specific gravity levels. All participants herein contained urine specific gravity values of ≤ 1.030 and were considered well hydrated.

### DXA/Body Composition

Participants’ height and weight was measured using a digital scale (Seca 769; Hanover), followed by body composition testing. Briefly, participants were asked to lay in the supine position for a minimum 5 minutes prior to full-body dual-energy x-ray absorptiometry (DXA) scan (Lunar Prodigy, GE Healthcare). Lean/soft tissue mass (also commonly termed as lean body mass; indicated as LBM herein) was assessed using associated software, and all scans were performed by the same technician who produced high test-retest reliability scores for LBM [intraclass correlation (ICC) = 0.99, standard error of the measurement (SEM) = 0.36 kg, and minimum difference (MD) value = 0.99 kg].

### Ultrasound

Ultrasonography images of the vastus lateralis (VL) were acquired with a B-mode imaging device (LOGIQ S7 Expert, GE Healthcare) using a multi-frequency linear array transducer (3-12 MHz, GE Healthcare). Participants were instructed to lay on an athletic training table in the supine position 5 minutes prior to imaging. Mid-thigh location was determined at 50% of total length between inguinal crease and proximal patella. Images were captured in the transverse plan using the B-mode function, settings were kept consistent across all participants, except for depth (Frequency: 12MHz, Gain: 55dB, Dynamic range: 72). Images were analyzed in ImageJ (National Institutes of Health) using either the polygon (VL muscle cross-sectional area, or VL mCSA) or segmented line (VL Thickness) functions. All images pre and post training were captured by the same technician who produced high test-retest reliability scores for VL thickness (ICC = 0.96, SEM = 0.09 cm, and MD = 0.24 cm) and VL mCSA (ICC = 0.99, SEM = 0.60 cm^2^, and MD = 1.65 cm^2^).

### Vastus lateralis muscle biopsies

Skeletal muscle biopsies were collected from the right vastus lateralis at the marked location from ultrasound imaging as previously described [31, 32]. Muscle samples were separated for DNA, RNA, and protein work (∼30-50 mg), placed in pre-labeled foil and flash-frozen in liquid nitrogen within two minutes of the biopsy. For immunohistochemistry (IHC; ∼20-30 mg), samples were mounted in Tissue-Tek® OCT Compound (Sakura Finetek Inc; cat #: 4583), slow frozen in liquid nitrogen-cooled isopentane, and subsequently stored at -80°C.

### Muscle tissue processing and analytical techniques

#### DNA Isolation

Frozen muscle samples (∼15 mg) were placed on a liquid nitrogen-cooled ceramic mortar, pulverized under liquid N2 and DNA was isolated using the DNeasy Blood and Tissue Kit (Qiagen; cat #: 69504) following manufacturer’s recommendations. DNA was eluted in buffer provided by the kit and DNA concentrations were determined in duplicate using a desktop spectrophotometer (NanoDrop Lite, Thermo Fisher Scientific).

#### RNA Isolation and targeted qPCR

Approximately 10 mg of tissue was placed in 500 µl of Trizol (Invitrogen; cat #: 15596026), and RNA isolation was conducted following manufacturer’s instructions. RNA concentrations were determined in duplicate using a NanoDrop Lite, followed by cDNA (2 µg) synthesis using a commercial qScript cDNA SuperMix (Quantabio; cat #: 95048-025).

Targeted real-time qPCR was performed in a thermal cycler (Bio-Rad Laboratories) using SYBR green (Quantabio; cat #: 95053-500) as routinely performed by our laboratory [33-35]. Gene-specific primers were designed using publicly available software (Primer3Plus), all primer sets and melt curves indicated only one amplicon was present. Forward and reverse primer sequences of all genes are listed in Table 1, and the rDNA primer set was designed per the report by Feng et al. [36] who used this set to quantify rDNA copy number in human tissue. Relative rDNA and rRNA values were acquired using the 2^ΔΔCq^ method, where 2^ΔCq^ = 2^(housekeeping gene (HKG) Cq - gene of interest Cq)^. GAPDH was used as the normalizer gene for all mRNA and rDNA data.

**Table 1.**
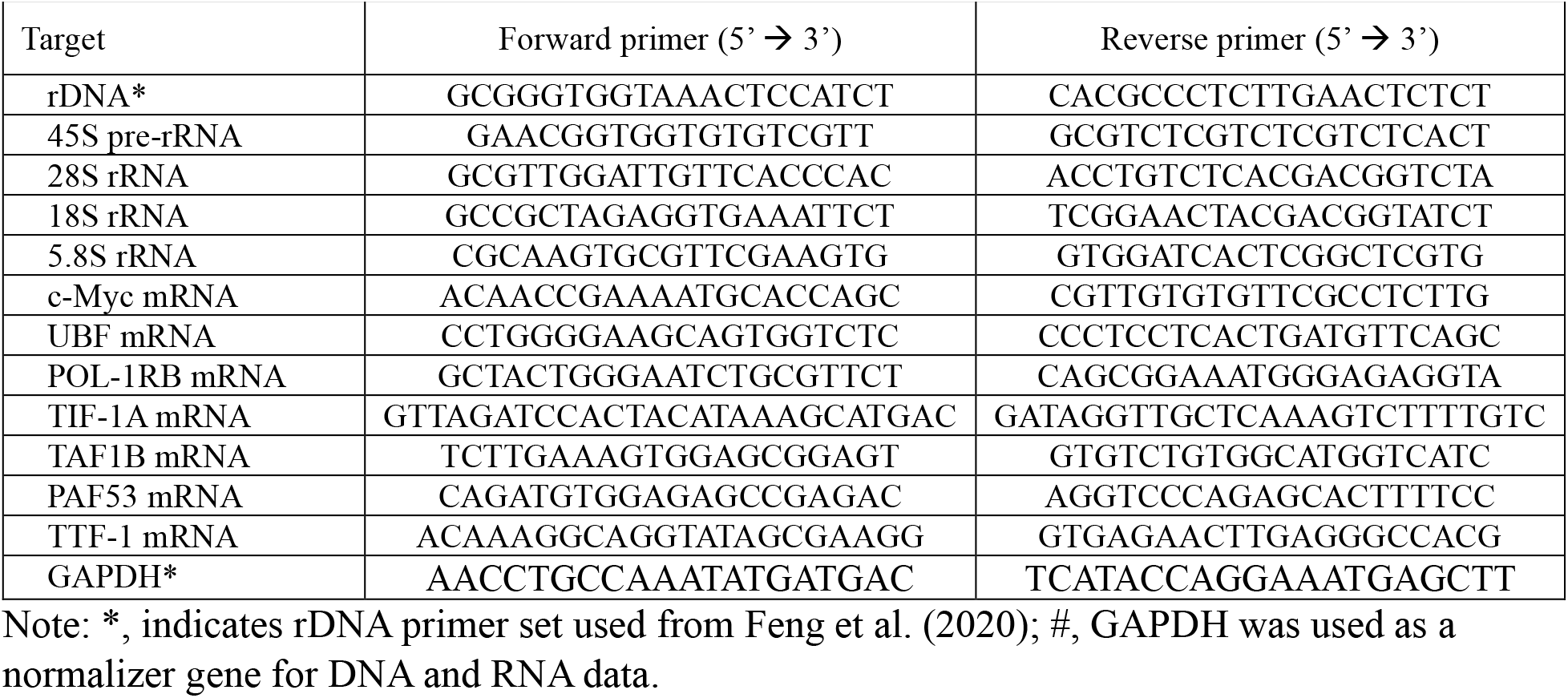
Human primer sets.

### Immunohistochemistry for fCSA

Although muscle sections were analyzed for fiber type-specific cross-sectional area (fCSA) as previously described [31, 32], only mean (weighted average of type I and II) fCSA data were used herein. Nonetheless, IHC methods are provided here for convenience to readers. Briefly, OCT-preserved samples were sectioned at a thickness of 10 µm using a cryotome (Leica Biosystems) and adhered to positively charged glass slides.

Sections were air dried for ∼30 minutes at room temperature followed by permeabilization using Triton-X (0.5%) in phosphate buffer solution (PBS) for 5 minutes. Following permeabilization, sections were washed for 5 minutes in PBS and immediately incubated in blocking solution for 30 minutes (Pierce Super Blocker, Thermo Fisher Scientific). Sections were then incubated in a primary antibody cocktail for 60 minutes containing 1x PBS, 5% Pierce Super Blocker Solution, 1:50 rabbit anti-dystrophin IgG1 (Genetex; cat #: GTX15277) and 1:50 mouse anti-myosin I IgG1 (Developmental Studies Hybridoma Bank; cat #: A4.951 supernatant). Following primary antibody cocktail incubation, sections were washed for 5 minutes in PBS and incubated with a secondary antibody cocktail for 60 minutes, containing 1x PBS, 1:100 anti-rabbit IgG1 conjugate Texas Red (Vector Laboratories; cat #: TI-1000), and 1:100 anti-mouse IgG1 conjugated AF488 (Thermo Fisher Scientific; cat #: A-11001). Sections were then washed for 5 minutes in PBS and mounted using a 4,6-diamidino-2-phenylindole containing fluorescent media (DAPI) (Genetex; cat #: GTX16206).

After staining, multiple digital images were captured using a fluorescence microscope (Nikon Instruments) with a 10x objective lens. Fiber-type specific fCSA was automatically analyzed for all images using open-sourced software, MyoVision [37].

### LHCN-M2 Immortalized Human Myotube Experiments

The LHCN-M2 human skeletal muscle cell line was kindly provided by Dr. Andrew Ludlow (University of Michigan, Ann Arbor, MI, USA). Cells were grown in 6-well plates at 37°C in 5% CO_2_ using growth media consisting of 4:1 of DMEM/M199 (Corning; Corning, NY, USA) supplemented with 15% fetal bovine serum (Avantor® Seradigm, VWR; cat #: 89510-182), 50 mM HEPES (Invitrogen; cat #: 15630-080), 0.03 µg/ml zinc sulfate (Thermo Fisher Scientific; cat # Z68-500), 1.4 µg/ml vitamin B12 (Sigma; cat #: V2876), 55 ng/ml dexamethasone (R&D Systems, distributed by VWR; VWR cat #: 1000505-728), 10 ng/ml human basic fibroblastic growth factor (BioPioneer; cat #: HRP-0011), and 2.5 ng/ml human hepatocyte growth factor (Chemicon International; cat #: GF116). Differentiation was induced at ∼85% confluency by switching to media containing 4:1 DMEM/M199 supplemented with 5% normal horse serum (Invitrogen; cat #: 26050-088). Following seven days of differentiation cells were treated with either: i) control media consisting of 0.1% DMSO (Corning; cat #: 25-950-CQC) for 24 hours, ii) 200 ng/ml IGF-1 (R&D Systems; cat #: 791-MG-050) for 24 hours, iii) 100 μM bisphenol-A (BPA; Sigma; cat #: 133027) for 24 hours, or iv) 100 μM BPA for 24 hours followed by 200 ng/ml of IGF-1 for 24 hours. Batch experiments contained 3-4 replicates per treatment condition.

Before the collection of the first batch of myotube lysates, 1 mM of puromycin dihydrochloride (VWR; cat#: 97064-280) was added to the media for assessment of muscle protein synthesis (MPS) using the surface sensing of translation (SUnSET) method [38]. Following pulse-labeling with puromycin 15 minutes prior to lysis, cells were lysed in 1 ml of Trizol reagent and placed in -20°C until further separation and isolation of DNA, RNA, and protein. Briefly, DNA, RNA, and protein were separated using the Trizol-bromochloropropane method [39, 40]. DNA was isolated using the commercially available DNeasy Blood and Tissue Kit (Qiagen; Germantown, MD, USA) and RNA was isolated using the commercially available Direct-zol RNA Miniprep Plus Kit (ZYMO; Irvine, CA, USA) following manufacturer’s recommendations. Protein was isolated as previously described by our laboratory [39], and quantification of protein isolates was determined using commercially available RC DC assay (Bio-Rad Laboratories; cat #: 5000121). Sample preps (1 µg/µl) were made to use in subsequent western blotting.

For imaging, myotubes were fixed in 4% PFA for 15 minutes at 37°C, washed three times in PBS/0.2% Triton-X for 3 minutes followed by a blocking step with 1%BSA/PBS/0.2% Triton-X for 1 hour at room temperature. Cells were then incubated with antibodies (1:100 in culture media) against sarcomeric myosin (Developmental Studies Hybridoma Bank; cat #: A4.1025) and embryonic myosin (Developmental Studies Hybridoma Bank; cat #: F1.652) for 3 hours at room temperature. Following three 3-minute washes, cells were incubated with goat anti-mouse IgG2a AF488 (ThermoFisher; cat#: A-21131) and goat anti-mouse IgG1 AF594 (ThermoFisher; cat#: A-21125) in PBS/0.2% Triton-X for 2 hours at room temperature. After incubation with secondary antibodies cells were washed, incubated with DAPI (ThermoFisher; cat#: D3571) for 10 minutes, and multiple images were obtained using a fluorescent microscope (Nikon Instruments) with a 20x objective lens.

### Statistics

Data were plotted and analyzed in GraphPad Prism (v10.1.0). Relative rDNA copy number (males and females), as well as rRNA and mRNA responses (females only), were examined for normality using Shapiro-Wilk tests. In females, rRNA and mRNA data were analyzed either using one-way repeated measures ANOVAs (normally distributed data) or Friedman tests (non-normally distributed data). Holm-Šídák’s multiple comparisons tests were used to decompose significant model effects for normally distributed data. Dunn’s multiple comparison tests were used to decompose significant model effects for non-normally distributed data. Relative rDNA copy number and rRNA/mRNA responses were associated with hypertrophic outcomes in males and/or females using Pearson correlations. One-way ANOVAs with Tukey’s post hoc tests were used to analyze *in vitro* outcomes. Statistical significance was established as p<0.05, and all data are expressed or plotted as mean and standard deviation values throughout.

## RESULTS

### rDNA copy number is not associated with the anabolic response or changes in ribosome content following 10-12 weeks of resistance training

Males experienced increases in LBM (+2.5±1.7 kg, p<0.001), VL thickness (+18.0±11.8%, p<0.001), mean fCSA (+15.9±24.3%, p<0.001), and total RNA (25.2±36.7%, p<0.001). Females (10 weeks of training, 2 d/week) experienced increases in LBM (+1.1±1.2 kg, p<0.001), VL thickness (+10.6±10.2%, p<0.001), and mean fCSA (+20.0±26.4%, p=0.002), though total RNA increases did not reach statistical significance (+11.4±43.6%, p=0.323). In females, mid-thigh VL muscle cross-sectional area (mCSA) increased with training (+11.8±12.0%, p<0.001); note, these assessments were not performed in the male participants.

Relative muscle rDNA copy number was not significantly different between the males and females (Fig. 1a). Relative rDNA copy number was not associated with training-induced changes in LBM, VL thickness, or mean fCSA (Fig. 1b-d), or changes in muscle RNA concentrations (Fig. 1e).

**Figure 1.**
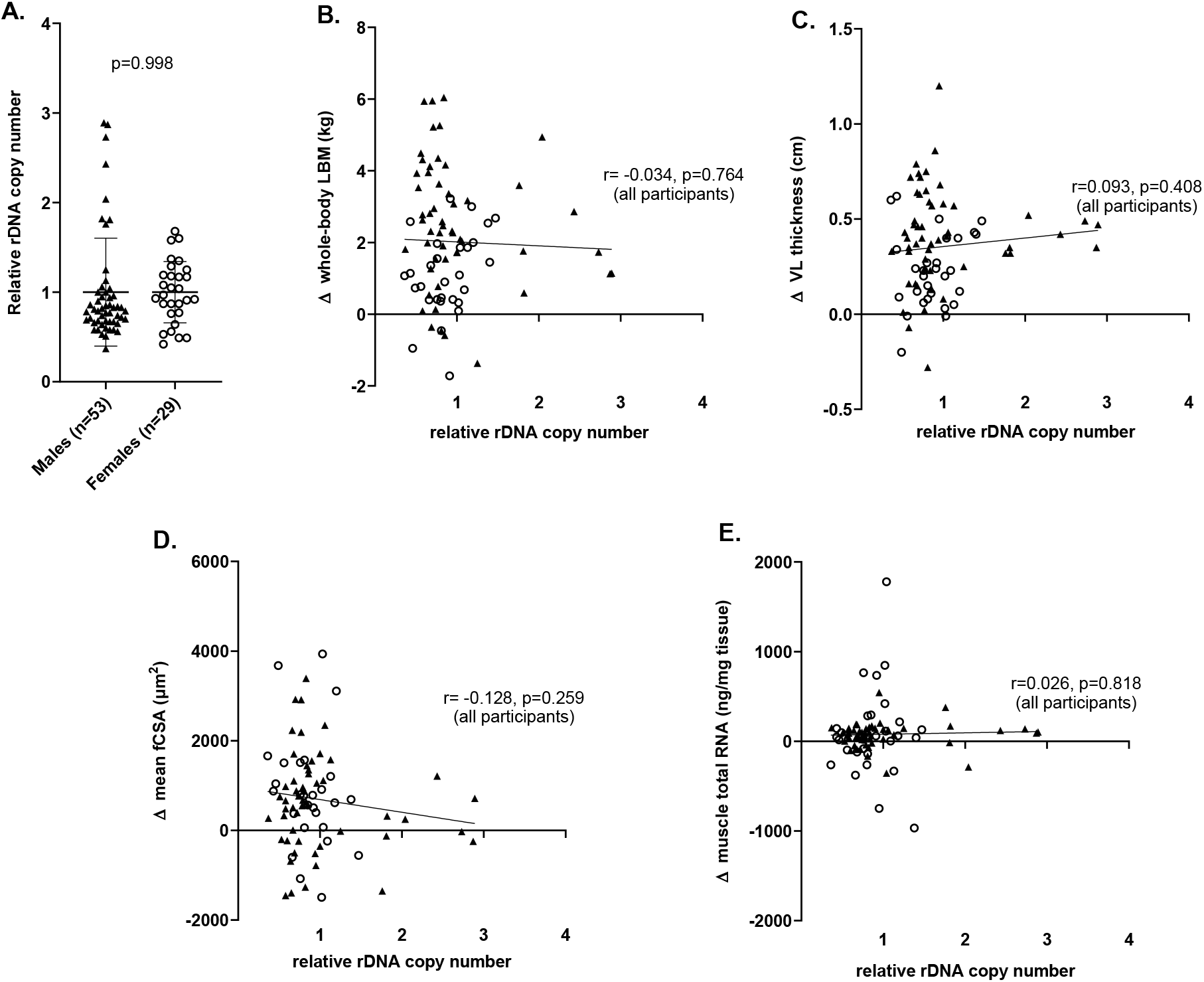
Associations between skeletal muscle relative rDNA copy number and skeletal muscle hypertrophic outcomes following 10-12 weeks of resistance training in all participants. Legend: Results from males (n=53) and females (n=29) who partook in 10-12 weeks of resistance training (2-3 days per week). Relative rDNA copy number was assessed from pre-training vastus lateralis (VL) muscle biopsies using qPCR (panel a). Associations were performed between relative rDNA copy number and the training-induced changes in whole-body lean body mass (LBM) assessed using dual-energy x-ray absorptiometry (panel b), VL thickness assessed using B-mode ultrasound (panel c), and VL mean (type I & II) myofiber cross-sectional area values (fCSA) using immunohistochemistry (panel d). An additional association was performed between relative rDNA copy number and training-induced changes muscle VL total RNA concentrations (i.e., ribosome content) (panel e).

### Markers of ribosome biogenesis are responsive to one bout and 10 weeks of resistance training in females

In addition to pre- and post-intervention biopsies, female participants underwent a biopsy 24 hours following their first resistance exercise (RE) bout. Changes in 45S pre-rRNA, 28S rRNA, and 18S rRNA were significantly elevated 24 hours following the first RE bout, and an elevation in 45S pre-rRNA was also observed following 10 weeks of training (Fig. 2a). A significant model effect was evident for 5.8S rRNA (p=0.025; Fig. 2a); however, post hoc tests indicated no significant differences between time points. c-Myc mRNA robustly increased 24 hours following the first RE bout (Fig. 2b), and UBF mRNA significantly increased following 10 weeks of training (Fig. 2c). All assayed components of the Pol-I regulon exhibited significant mRNA elevations acutely (e.g. 24 hours following the first RE bout) (Fig. 2d).

**Figure 2.**
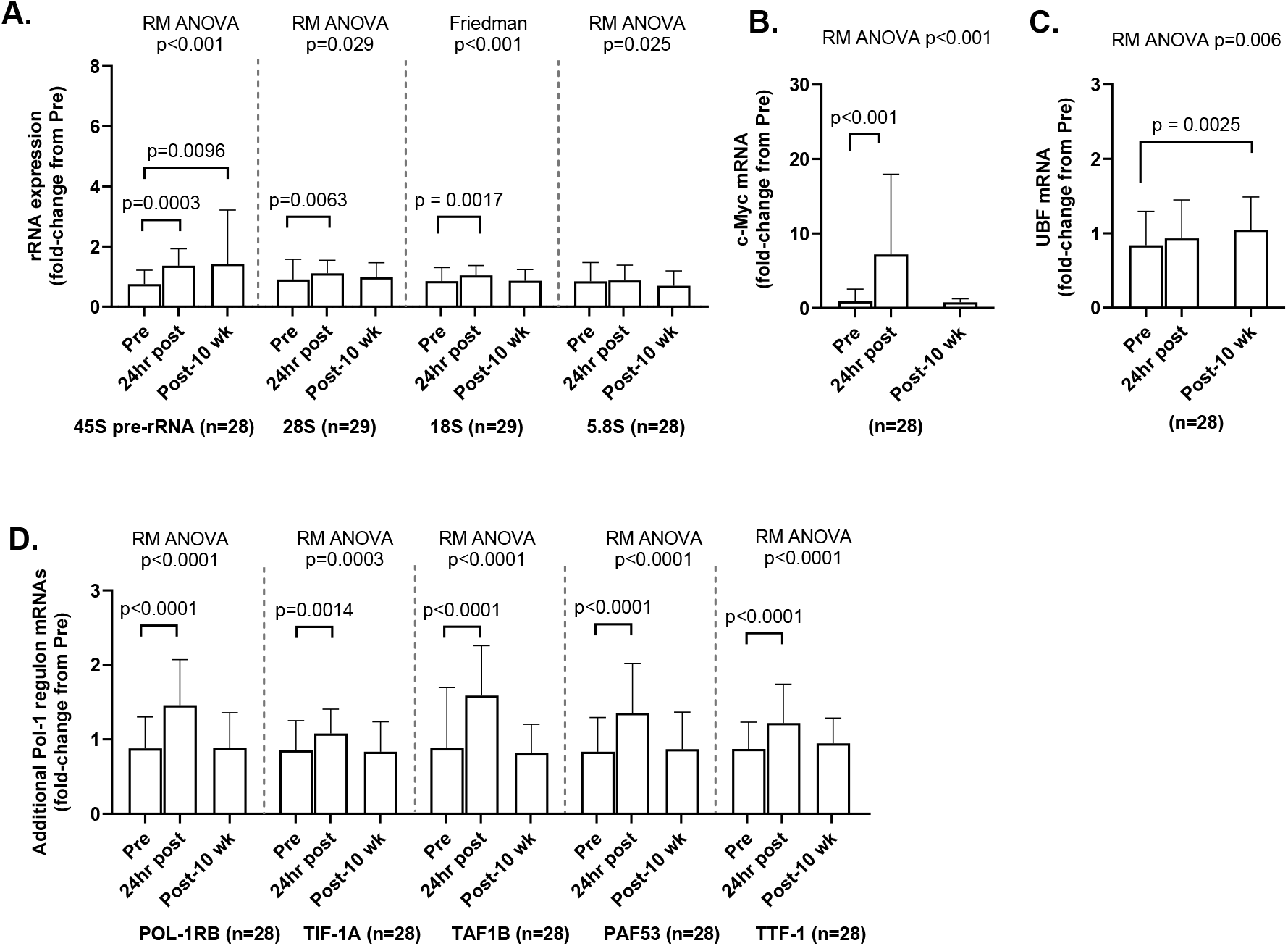
Alterations in markers of ribosome content and ribosome biogenesis following one bout and 10 weeks of resistance training in females. Legend: Results from females who partook in 10 weeks of resistance training (2 days per week). Unlike the males in Fig. 1, a biopsy was taken 24 hours following the first bout of training in these participants (noted as 24hr post). Different rRNA transcripts were assessed using qPCR (panel a), mRNAs of transcriptional regulators of ribosome biogenesis are presented (panels b/c), and additional RNA polymerase I Pol-I regulon mRNAs are presented (panel d). Bar graphs are presented as means with positive standard deviation bars and n-sizes per variable are noted in each panel. Significance from repeated measures (RM) ANOVAs, Friedman tests, and post hoc tests are also noted accordingly.

### rDNA copy number is not associated with rDNA transcription following a bout of resistance exercise in females

Given that certain rRNA transcripts (i.e., 45S pre-rRNA, 28S rRNA, and 18S rRNA) were elevated 24 hours following the first RE bout in females, we examined these changes relative to rDNA copy number. As shown in Figure 3 (panels a-d), none of these transcripts significantly correlated with rDNA copy number.

**Figure 3.**
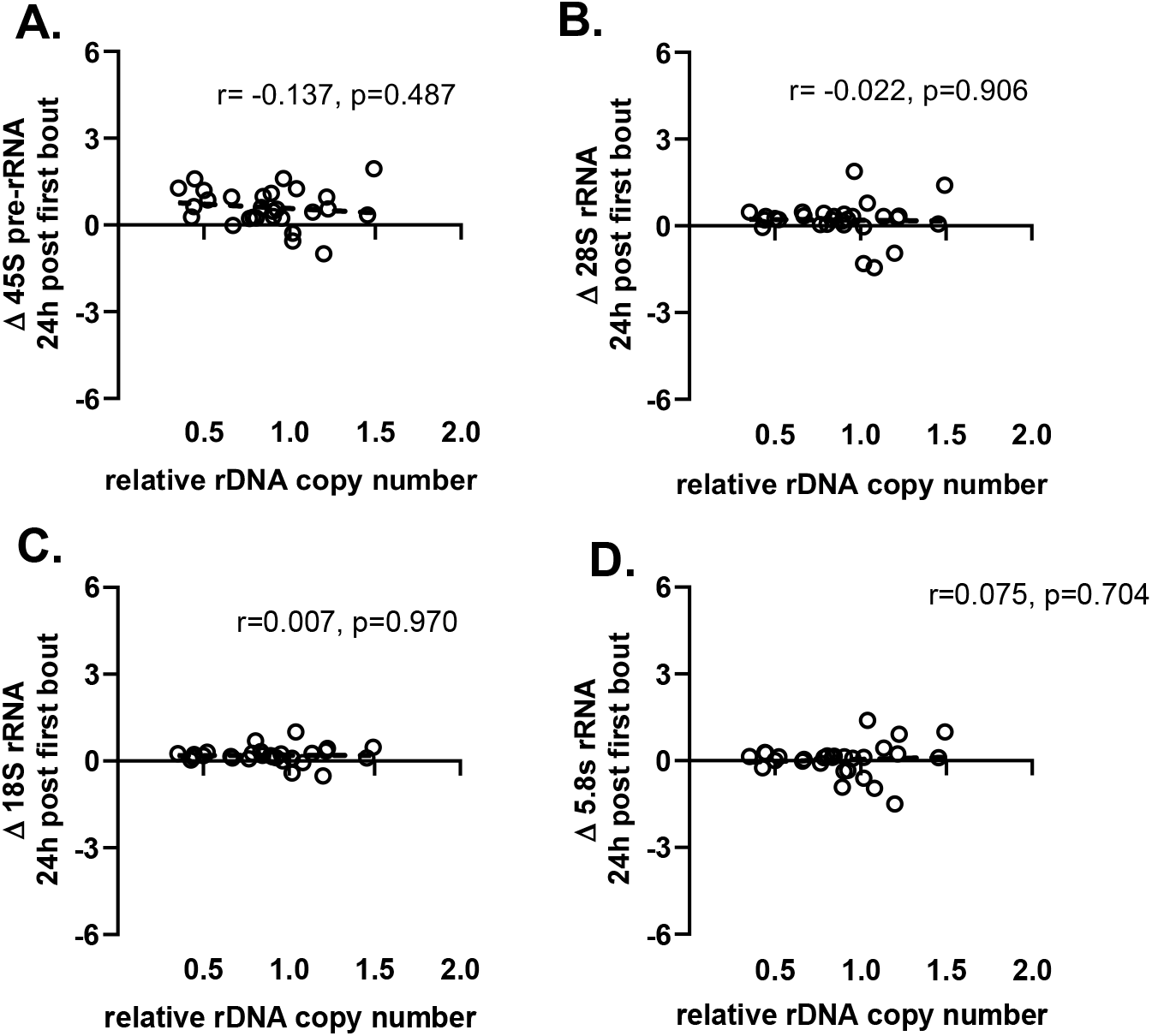
Female-only associations between rRNA responses 24 hours following the first workout bout and relative rDNA copy number in skeletal muscle. Legend: Results demonstrating lack of associations between relative rDNA copy number and the acute (Pre→ 24-hour post bout 1) transcript responses for 45S pre-rRNA (panel a), 28S rRNA (panel b), 18S rRNA (panel c), and 5.8S rRNA. Note: all variables contain data from 28 females except 45S pre-rRNA which contains data from 27 females.

### Elevations of select ribosome biogenesis transcripts following acute resistance exercise correlates with hypertrophic outcomes following 10 weeks of training

Table 2 contains select associations in female participants. As shown in Figure 1, relative rDNA copy number was not associated with hypertrophic outcomes in all 82 participants (panels b-d). The top portion of Table 2 contains Pearson correlation values of relative rDNA copy number and hypertrophic outcomes in the females only, and again, no significant associations were evident. However, significant positive associations were evident between 10-week changes in VL mCSA and the acute (Pre → 24-h following first RE bout) fold-changes in two of the 10 assayed transcripts related to rDNA transcription (28S rRNA and c-Myc mRNA). Additionally, significant positive associations were evident between 10-week changes in mean fCSA and the acute (Pre**→** 24-h post bout 1) changes in four of the 10 assayed mRNAs related to rDNA transcription (UBF, POL-1RB, TAF1B, and PAF53).

**Table 2.**
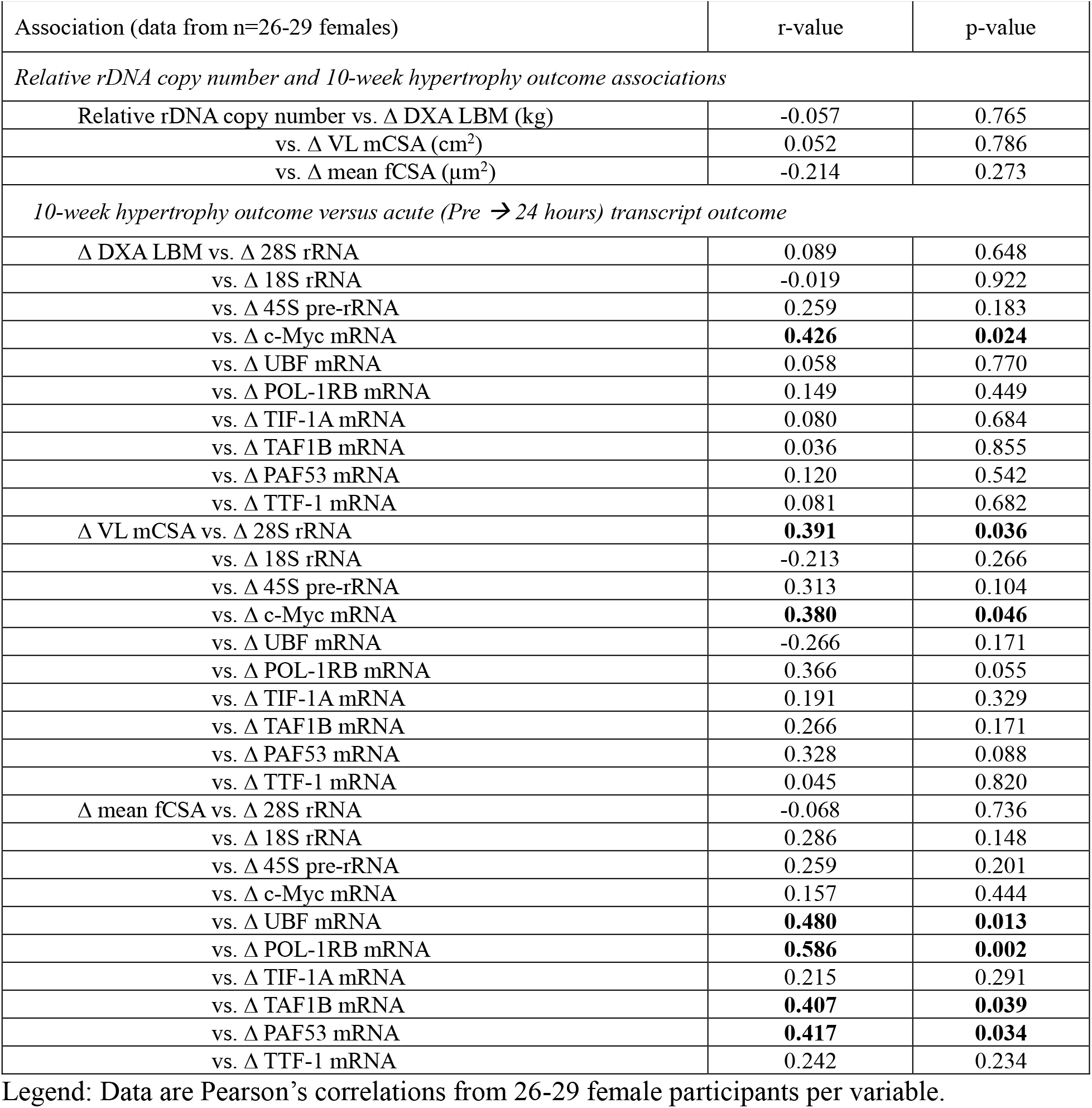
Acute bout transcript and hypertrophy associations in female participants only.

### Elevations of select ribosome biogenesis transcripts following resistance training correlate with training-induced hypertrophic outcomes

Table 3 contains associations between pre-to-post intervention fold-changes in ribosome biogenesis transcripts and changes in hypertrophic outcomes. Notably, significant positive associations were evident between 10-week changes in mean fCSA and fold-changes in four of the 10 transcripts assayed related to rDNA transcription (45S rRNA and mRNAs for UBF, POL-1RB, and PAF53).

**Table 3.** Chronic training transcript and hypertrophy associations in female participants only.

| Association (data from n=26-29 females) | r-value | p-value |
| --- | --- | --- |
| *10-week hypertrophy outcome versus chronic (Pre → Post) transcript outcome* |  |  |
| $\Delta$ DXA LBM vs. $\Delta$ 28S rRNA | 0.108 | 0.578 |
| vs. $\Delta$ 18S rRNA | 0.141 | 0.466 |
| vs. $\Delta$ 45S pre-rRNA | 0.152 | 0.440 |
| vs. $\Delta$ c-Myc mRNA | 0.143 | 0.467 |
| vs. $\Delta$ UBF mRNA | 0.112 | 0.571 |
| vs. $\Delta$ POL-1RB mRNA | 0.205 | 0.296 |
| vs. $\Delta$ TIF-1A mRNA | 0.140 | 0.478 |
| vs. $\Delta$ TAF1B mRNA | 0.091 | 0.647 |
| vs. $\Delta$ PAF53 mRNA | 0.207 | 0.290 |
| vs. $\Delta$ TTF-1 mRNA | **0.425** | **0.024** |
| $\Delta$ VL mCSA vs. $\Delta$ 28S rRNA | **0.373** | **0.046** |
| vs. $\Delta$ 18S rRNA | **0.531** | **0.003** |
| vs. $\Delta$ 45S pre-rRNA | -0.048 | 0.810 |
| vs. $\Delta$ c-Myc mRNA | -0.024 | 0.902 |
| vs. $\Delta$ UBF mRNA | 0.150 | 0.446 |
| vs. $\Delta$ POL-1RB mRNA | 0.145 | 0.463 |
| vs. $\Delta$ TIF-1A mRNA | 0.057 | 0.774 |
| vs. $\Delta$ TAF1B mRNA | 0.030 | 0.880 |
| vs. $\Delta$ PAF53 mRNA | 0.263 | 0.176 |
| vs. $\Delta$ TTF-1 mRNA | 0.208 | 0.288 |
| $\Delta$ mean fCSA vs. $\Delta$ 28S rRNA | 0.062 | 0.757 |
| vs. $\Delta$ 18S rRNA | 0.123 | 0.540 |
| vs. $\Delta$ 45S pre-rRNA | **0.478** | **0.011** |
| vs. $\Delta$ c-Myc mRNA | 0.046 | 0.825 |
| vs. $\Delta$ UBF mRNA | **0.400** | **0.044** |
| vs. $\Delta$ POL-1RB mRNA | **0.460** | **0.018** |
| vs. $\Delta$ TIF-1A mRNA | 0.199 | 0.330 |
| vs. $\Delta$ TAF1B mRNA | 0.189 | 0.354 |
| vs. $\Delta$ PAF53 mRNA | **0.474** | **0.014** |
| vs. $\Delta$ TTF-1 mRNA | 0.280 | 0.166 |

Legend: Data are Pearson's correlations from 26-28 females per variable.

### Reductions in rDNA copy number do not affect myotube anabolism

To validate the lack of association between relative rDNA copy number and hypertrophic outcomes in humans, we designed mechanistic experiments to determine whether reducing rDNA copy number affected myotube anabolism *in vitro*. Bisphenol A (BPA, 100 µM) was used to reduce rDNA copy number [30], and insulin-like growth factor-1 (IGF1, 200 ng/ml) was used to induce anabolism [41]. Exposure to BPA for 24 hours caused a significant reduction in rDNA copy number (Fig. 4a). Reducing rDNA copy number did not reduce myotube diameter and did not prevent IGF1 induced cellular hypertrophy (Fig 4b). Consistently, BPA did not affect basal protein synthesis nor altered the anabolic response to IGF1 (Fig 4d). BPA and/or IGF1 treatments did not significantly affect rRNA concentrations (Fig. 4f-h) or 45S pre-rRNA concentrations (Fig. 4i). In line with our human associations (Fig. 3), we interpret these data to suggest that a reduction in relative rDNA copy number does not interfere with rDNA transcription *in vitro*.

**Figure 4.**
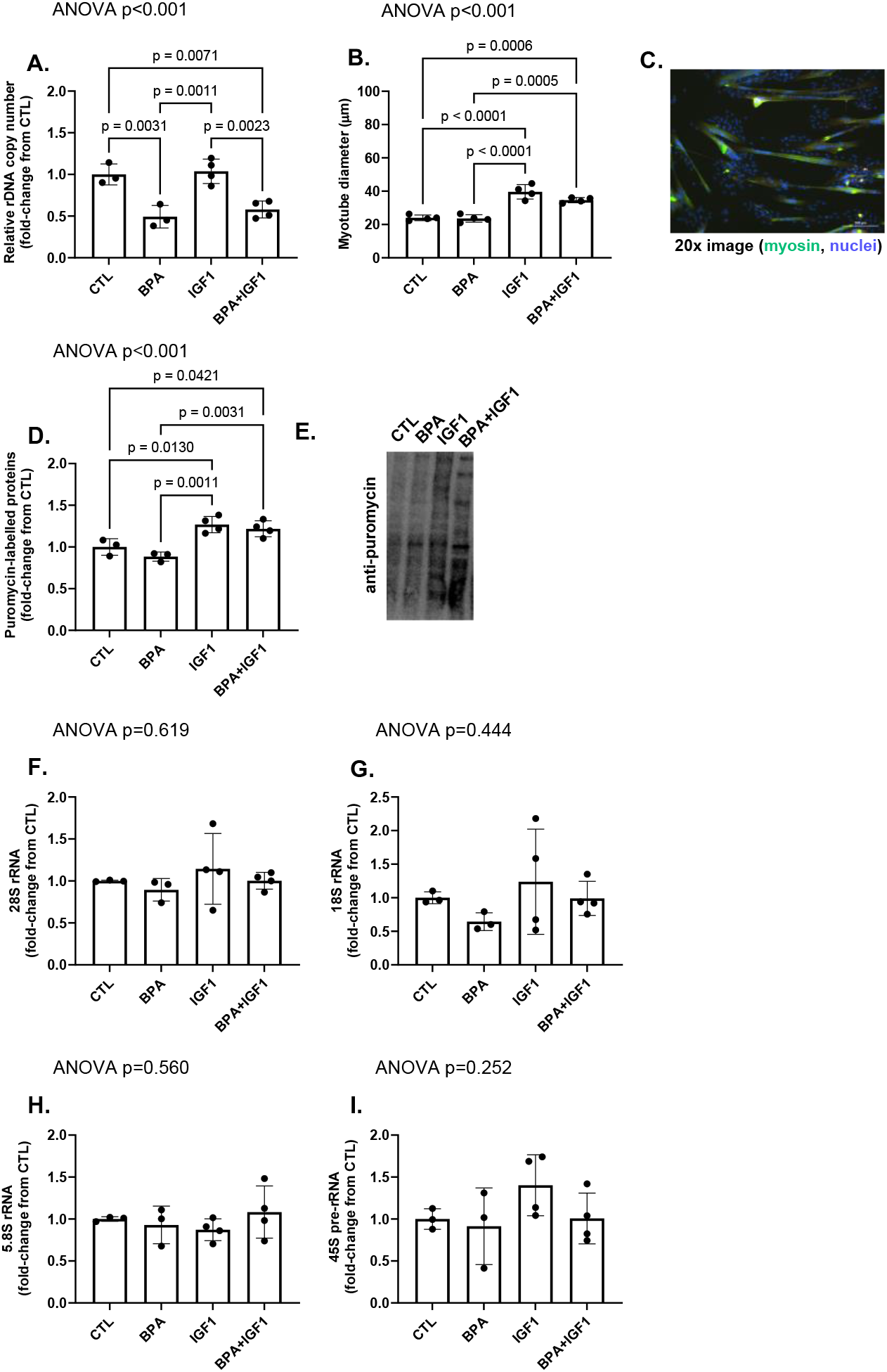
Effects of rDNA copy number reduction on the anabolic response to IGF1 in human myotubes. Legend: The effects of 24-hour bisphenol A (BPA, 100 µM) and/or insulin-like growth factor-1 (IGF1 200 ng/ml) treatments on relative rDNA copy number (panel a), myotube diameter (panel b), protein synthesis (panel d), and relative rRNA content (panels F-I). Treatments consisted of 3-4 replicates, data were analyzed between treatments using one-way ANOVAs and Tukey’s post hoc tests, and data are graphed as individual values with mean ± standard deviation bars. Images: panel c, representative image used for myotube diameter analysis; panel e, representative puromycin western blot. Other note: CTL, control (untreated) cells.

## DISCUSSION

Increases in ribosomal content and ribosome biogenesis are well characterized following resistance training bouts. The current data add to the literature by demonstrating a robust increase in ribosomal content (rRNAs) and various ribosome biogenesis related markers, namely Pol-I regulon mRNAs, 24 hours following a resistance exercise bout in untrained females.

Additionally, UBF and c-Myc mRNAs, along with the mRNA expression of other Pol-I regulon factors, were upregulated in females following resistance training. Critically, each of these targets contribute to an increase in rDNA transcription [10, 14], and we demonstrated that increases in numerous ribosome content and ribosome biogenesis related markers were associated with increases in skeletal muscle fiber hypertrophy.

The novel and primary finding herein are the lack of associations between relative rDNA copy number and training-induced changes in skeletal muscle hypertrophy outcomes among 82 participants. Relative rDNA copy number was also not significantly associated with 24-hour post exercise ribosome biogenesis markers in females. It has been posited that a relatively higher rDNA copy number may contribute to the hypertrophic response to mechanical overload by allowing for a larger increase in ribosome biogenesis [1, 3, 42-44]. In line with this hypothesis, Figueiredo et al. [16] reported a significant association exists between relative rDNA copy number and 24-hour post-resistance exercise 45S rRNA expression in humans. Our findings do not align with this previous report. On possible explanation for this discrepancy is the difference in the number of participants between studies. Figueiredo et al. examined this relationship in 7 male and female participants, whereas our 24-hour post-exercise correlation data included 29 females. Hence, the larger sample size (as well as participants being female-only) in our study likely accounts for disparate findings. Given that this is the first study to examine relative rDNA copy number and the hypertrophic response to resistance training, no comparison to prior data is possible. Hence, the novel finding of relative rDNA copy number not being associated with mechanical overload-induced skeletal muscle hypertrophy warrants confirmation. However, the associations between fold-changes in ribosome biogenesis markers and resistance training-induced skeletal muscle hypertrophy agrees with prior literature [19, 21, 45].

We provide further support for our interpretation that rDNA copy number does not affect muscle hypertrophy. Using BPA to reduce rDNA copy number of human-derived primary myotubes [30], we demonstrate that a reduction in rDNA copy number did not affect myotube diameter, protein synthesis, or markers of ribosome biogenesis. Importantly, IGF-1 treatment of myotubes with reduced rDNA copy number resulted in similar anabolic responses compared to untreated myotubes with lower rDNA copies. Again, these findings suggest that protein synthesis and rDNA transcription (irrespective of the relative rDNA copy number) are more likely contributors to myofiber hypertrophy.

### Limitations

The current study has limitations, namely relative rDNA copy number was measured via qPCR as compared to whole-genome sequencing which would likely provide a more accurate measure of rDNA copy number. Additionally, all participants were younger, healthy individuals and the study design for the 53 male participants contained only pre- and post-training intervention time points, thus limiting the acute response analyses to females. Hence, further acute studies in males whereby relative rDNA copy number is examined in conjunction with acute ribosome biogenesis responses will strengthen the current conclusions.

## Conclusions

We provide evidence from both our larger human cohort and *in vitro* experiments demonstrating that relative rDNA copy number does not influence the hypertrophic outcome to resistance training or IGF-1 stimulation, respectively. Our results agree with a body of current literature supporting that rDNA transcription, amongst other mechanisms such as enhanced protein synthesis and myonuclear accretion, is a driver of skeletal muscle hypertrophy.

## AUTHOR CONTRIBUTIONS

J.S.G., G.A.N, and M.D.R. conceived the idea for this study. A.T.L., A.D.F., and M.D.R. provided significant resources to complete various project aims. J.M.M. performed multiple experiments throughout, which warranted co-authorship. J.S.G. and M.D.R. primarily drafted the manuscript, and all co-authors provided feedback as well as intellectual contributions. All authors have read and approved the final version of this manuscript and agree to be accountable for all aspects of the work in ensuring that questions related to the accuracy or integrity of any part of the work are appropriately investigated and resolved. All persons designated as authors qualify for authorship.

## ACKNOWLEDGMENTS

Costs for the human study and reagents were provided by the Peanut Institute (Albany, GA, USA) obtained by A.D.F., M.D.R, and others as well as discretionary lab funds of M.D.R. and A.D.F. The authors would like to thank Kaelin Young, Cody Haun, Paul Roberson, Petey Mumford, Matthew Romero, Bradley Ruple, Morgan Smith, Kristen Smith, Shelby Osburn, and previous lab members who critically assisted with the human interventions.

## DATA AVAILABILITY STATEMENT

The data that support the findings of this study are available from the corresponding author upon reasonable request.

